# Feeding secondary fermentations with mammalian and fungal culture waste streams increases productivity and resource efficiency

**DOI:** 10.1101/2023.10.17.562659

**Authors:** Ciara Lynch, Federico Cerrone, Kevin E O’Connor, David J O’Connell

**Author notes:** Federico Cerrone and David O’Connell are corresponding authors. **Email:**. **Author Contributions:** CL and FC collected experimental data, CL, FC, and DO’C wrote text, and CL, FC, DO’C and KO’C were responsible for editing. **Competing Interest Statement:** No competing interests to be disclosed.

## Abstract

The evolution of the circular bioeconomy will require the realisation of new value from waste streams generated from all manufacturing processes. Bioprocessing of recombinant proteins and functional foods using animal and microbial fermentations are fast growing industries with increasing volumes of spent culture media waste resulting in significant resource inefficiency. Here Chinese hamster ovary cell spent media (CSM) and *Trametes versicolor* fungal cell spent media (FSM) were used as model waste streams to feed *Escherichia coli* in secondary fermentations. Expression of a recombinant fusion protein was used to measure the value of these streams as feedstuff. *E. coli* cultures fed with either waste stream in bioreactors produced equivalent yields of the protein as those fed with rich microbiological media. Separately CSM was tested as a replacement feed for the fungal cell fermentation producing beta and alpha glucans. This waste-fed fermentation resulted in a 2-fold higher glucan yield than when using standard corn steep liquor feed. Quantitative analysis of the spent media content with mass spectrometry and hplc methods showed significant differences in elemental composition profiles, quantities of individual amino acids and carboxylic acids and levels of carbon sources. These findings underline the versatility of *E.coli* in utilisation of waste media as a feedstuff and show that the range of applications of CHO cell culture media waste are not limited to feeding bacterial fermentation. Exploration of cellular bioprocesses that can efficiently use cell media waste as fermentation feed will be a valuable step towards increased resource efficiency in bioprocessing industries. (249/250).

**Significance Statement:** The expanding range and scale of valuable biomolecular products developed with bioprocessing activities using animal and microbial cells is responsible for the production of hundreds of millions of litres of waste culture media annually. Recycling this waste stream has the potential to decrease the impact of the associated high water and energy consumption contributing to meeting targets for the Paris 2050 Agreement and UN Sustainable Development Goals. This work shows that waste media from distinct bioprocesses can be reused to sustain high-level recombinant protein production in a secondary fermentation and the notable finding that a waste media can add value to a secondary process by increasing its productivity. Efficient valorisation of waste media is an urgent and scientifically valuable objective. (120/120)

## Introduction

The implementation of a circular bioeconomy is a major goal in the attempt to bring humans back to living within the earth’s planetary boundaries, reducing the impact of human activities on climate change. Current research seeks to maximise resource efficiency by maintaining the value and utility of resources for as long as possible e.g., the reuse of waste within a process or use as a starting material for a secondary process (Stegmann, Londo and Junginger, 2020). This is reflected in a broad array of studies into the conversion of waste to products of value in the last ten years, from generation of valuable biomass to synthesis of bioplastics (Gao *et al*., 2016; Narancic *et al*., 2018, 2020; Diboune *et al*., 2019; Ruiz *et al*., 2019). One example of a waste intensive process yet to be assessed for conversion to products of value is the cell culture waste produced by the bioprocessing industry. There are currently 443 individual licensed biopharmaceutical products from vaccines to monoclonal antibodies that are made in cell culture bioprocess using expression hosts ranging from mammalian cells to microbes. Monoclonal antibodies, with over 80% of the total protein-based biopharmaceutical global sales in 2021, are principally made using Chinese hamster ovary cells (CHO) in intensified bioprocesses using chemically defined media at scales from hundreds to tens of thousands of litres (Walsh and Walsh, 2022). Global bioprocessing capacity has rapidly increased to provide millions of litres in production facilities worldwide to support demand for these molecules at tonnes scale. Currently in all of these facilities once the product is harvested the complex media formulation used to grow these cells is immediately sent for treatment as waste for disposal.

A newer area of cell culture bioprocessing predicted to undergo rapid growth is that of precision fermentation of synthetic meat and dairy products, designed to replace conventional foods from agricultural production (Lübeck and Lübeck, 2022; Boukid *et al*., 2023; Dupuis *et al*., 2023). Cultured meat using animal stem cells grown in bioreactors for example, has potential benefits proposed to lower greenhouse gas production from animals, reduce land use for animal farming, and reduce energy and water consumption helping to meet those targets for the Paris 2050 Agreement and UN Sustainable Development Goals (Post, 2012). However, to achieve a scale to meet global demand significant volumes of new bioprocessing waste will be generated that could dwarf those produced by the biopharmaceutical industry in litres per annum.

A number of pilot studies have investigated the principle of recycling spent media waste and show utility when used as a supplement to fresh media in the same bioreactor across a number of cell types and processes (Kempken, Büntemeyer and Lehmann, 1991; Hsiao, Glatz and Glatz, 1994; Riese *et al*., 1994; Wu, Ruan and Lam, 1998). We have previously shown spent media from CHO cell culture supports growth and recombinant protein production in a secondary fermentation as a standalone feed (Lynch and O’Connell, 2022). The nutritional value remaining in spent culture media could potentially be employed to feed multiple, distinct secondary fermentations. CHO cell spent culture media and spent culture media from *Trametes versicolor* fermentation represent models of biopharmaceutical bioprocess and precision fermentation of functional food ingredient waste streams. Filamentous fungi such as *T. versicolor* accumulate polysaccharides such as alpha- and beta-glucan in their cell walls (Lemieszek and Rzeski, 2012; Maehara *et al*., 2012; Kyanko *et al*., 2013). These molecules have demonstrated immunostimulant activities and are involved in the regulation of the slow absorption of glucose by gut associated lymphoid tissue (GALT) associated microbiota with health benefits against insulin resistance (Chan, Chan and Sze, 2009; Vetvicka, Novak and Silva, 2018; Gürünlü, 2022; Kozarski *et al*., 2023). Submerged cultivation of filamentous fungi is a recently devleoped bioprocess strategy to increase the yield of fungal biomass containing bioactive compounds, in comparison to the traditional and unsustainable compost-based methods of fungal production (Gibbs, Seviour and Schmid, 2000). A comparable amount of biomass is obtained in a matter of days in this way rather than weeks in the latter.

In this study we investigated production of a recombinant protein by *E.coli* in secondary fermentation in bioreactors fed with rich microbiological media or waste media from the model streams and compared the overall yields of recombinant protein in each. We also compared total glucan production from a *T. versicolor* secondary fermentation fed with CHO spent media with the yield from the standard fermentation feed. To understand the impact of waste media on each of these bioprocesses we then employed elemental composition analysis with ICP-MS and metabolite composition analysis with LC-MS/MS. Proteomic adaptations made by the *E. coli* and *T. versicolor* cells illuminate the versatility of expression hosts in successfully utilising these waste media feeds.

## Results

### Spent media from CHO and *Trametes versicolor* cell culture as nutrients in an *E. coli* fermentation

CHO spent media (CSM) and fungal spent media (FSM) were compared with respect to the growth rate of the *E*.*coli* culture and the recombinant protein expression levels (mCherry-EF2) in shake flask culture. *E.coli* cultures grown in the standard rich media of LB grew fastest with a specific k of 1.042, while *E.coli* grown in CSM achieved a growth rate (k) at ∼93% of LB grown cultures. FSM grown cultures had the lowest growth rate (∼88% of LB grown cultures) (Table 1). Recombinant protein yield of the CSM reached 68% of the LB fed cultures, while FSM reached 44% (Table 1).

**Table 1.** Growth rates and recombinant protein yields for flask and bioreactor experiments. Standard deviation (SD) measurements for flask experiments had n = 3, for bioreactors n = 2. Original data values used for calculations available in table S1.

| Measurement | LB | CSM | FSM |
| --- | --- | --- | --- |
| Flasks |  |  |  |
| Specific growth rate (k) | 1.042 | 0.928 | 0.884 |
| Doubling time (hrs) | 0.665 | 0.747 | 0.784 |
| R-squared coefficient | 0.998 | 0.995 | 0.996 |
| Protein produced (g/L ± SD) | 0.489 ± 0.017 | 0.333 ± 0.002 | 0.216 ± 0.008 |
| Bioreactors |  |  |  |
| Specific growth rate (k) | 0.589 | 0.264 | 0.454 |
| Doubling time (hrs) | 1.177 | 2.630 | 1.528 |
| R-squared coefficient | 0.991 | 0.951 | 0.950 |
| Cell dry weight (g/L) | 7.100 | 6.100 | 5.400 |
| Protein produced (g/L ± SD) | 1.292 ± 0.093 | 1.353 ± 0.076 | 1.190 ± 0.057 |

Comparison of the cultures in 0.5 L reactors for 20-hour expressions showed the growth of the *E. coli* in the CSM and FSM conditions remained slower than the control LB rich medium while conversely the recombinant protein yield was greater than or equal to the LB control with 1.35 g/L (CSM) and 1.19 g/L (FSM) compared to 1.14 g/L (LB) (Table 1).

Glycerol consumption was negligible across the conditions, except after induction of expression in the CSM condition, where a further ∼10 g/L was consumed by the end of the culture time frame (Fig 2). Detection of carbon sources by HPLC revealed that, while no glucose was supplemented to any media, the waste media contained glucose (9.08 and 2.92 g/L respectively) prior to use in secondary fermentation. LB on the other hand contained no glucose prior to starter culture addition. After addition of the starter culture, the glucose concentrations of the LB, CSM and FSM were 2.76, 9.77, and 5.14 g/L respectively. All glucose was fully consumed by the time expression was induced by IPTG addition (Fig 2).

**Figure 1.**
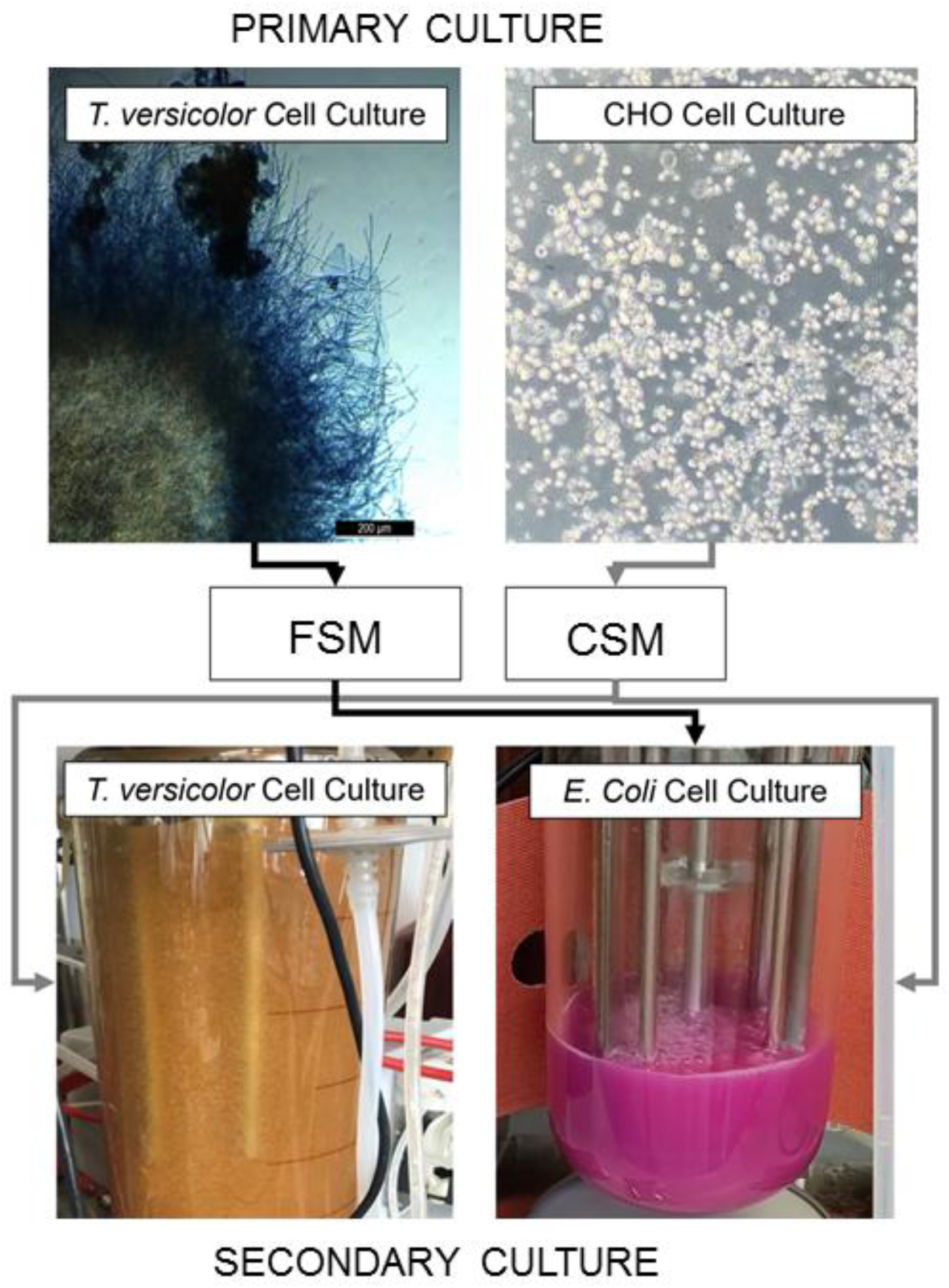
Flow diagram of spent media crossover between the two existing processes.

**Figure 2:**
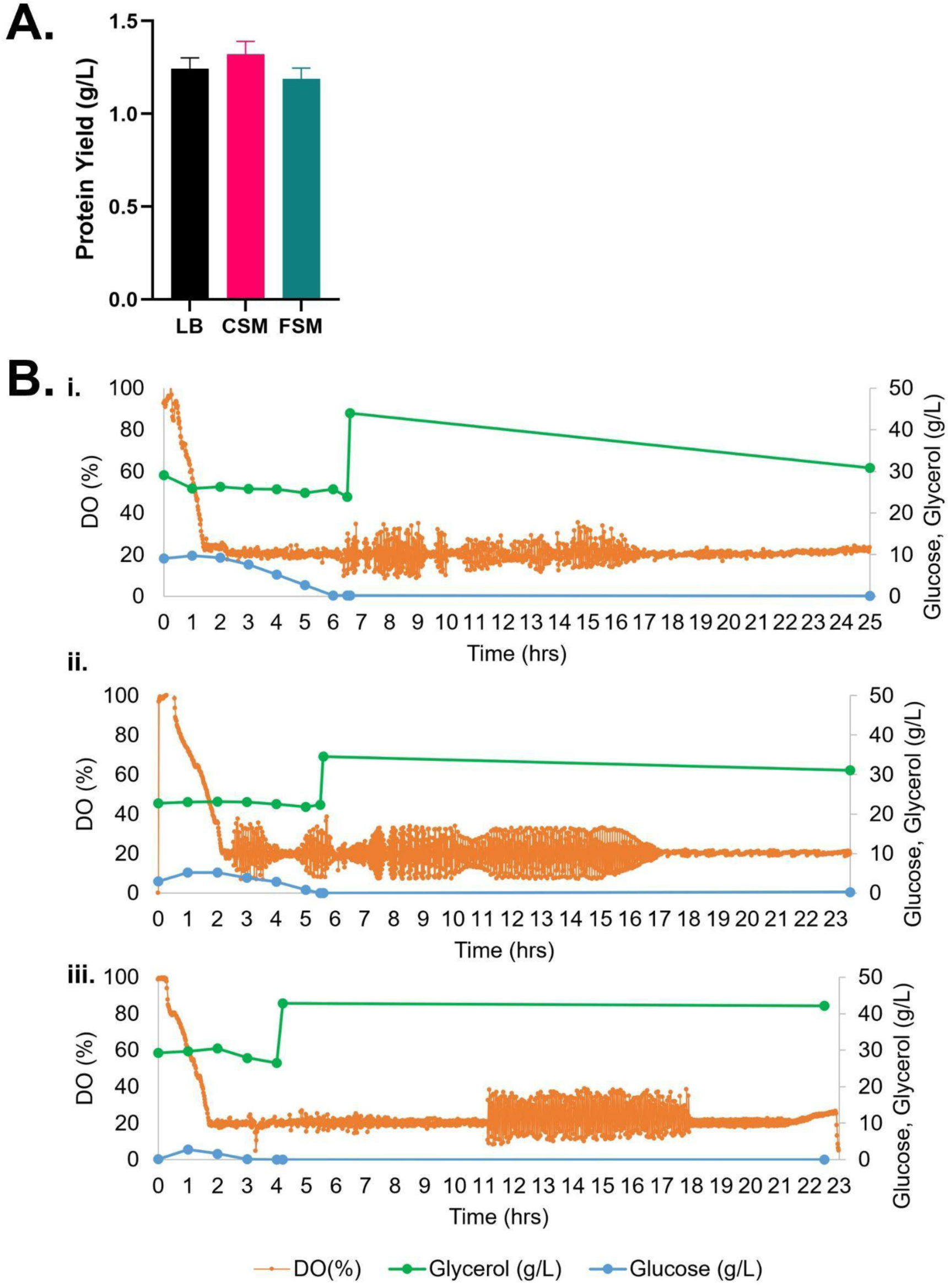
**A:** Recombinant protein yield of mCherry-EF2 from E. coli cultured in various media in bioreactors, LB = lysogeny broth, CSM = CHO spent media, FSM = fungal spent media. Error bars represent standard deviation, where n = 2. No statistical difference was seen in the conditions. **B:** Bioreactor timelapse of dissolved oxygen (DO) and carbon source consumption measured by HPLC (glycerol and glucose), i: CSM, ii: FSM, iii: LB. IPTG induction of protein expression was carried out along with addition of 2% glycerol once cultures reached OD_600 nm_ of 10. For raw data see supplementary table S1.

### Spent media from CHO cell culture as nutrient addition in secondary *Trametes versicolor* fermentation

The CSM was used to supplement a fungal media composed of 10 g/L of glucose and 3.3 g/L of KH_2_PO_4_. This was added as a 15 mL/L sterile solution to shake flask cultures of *Trametes versicolor*. This spent media-supplemented culture achieved an average CDW of 7.23 ± 0.49 g/L; this value is 13% higher than cultures grown in the typical corn steep liquor (CSL) control media (Table 2). Glucose was completely consumed after 8 days (the first 15% was consumed within 3 days of growth) and the total protein was reduced by 66%. 20% of the cell dry weight of *T. versicolor* biomass was composed of glucans after 8 days of growth.

**Table 2.**
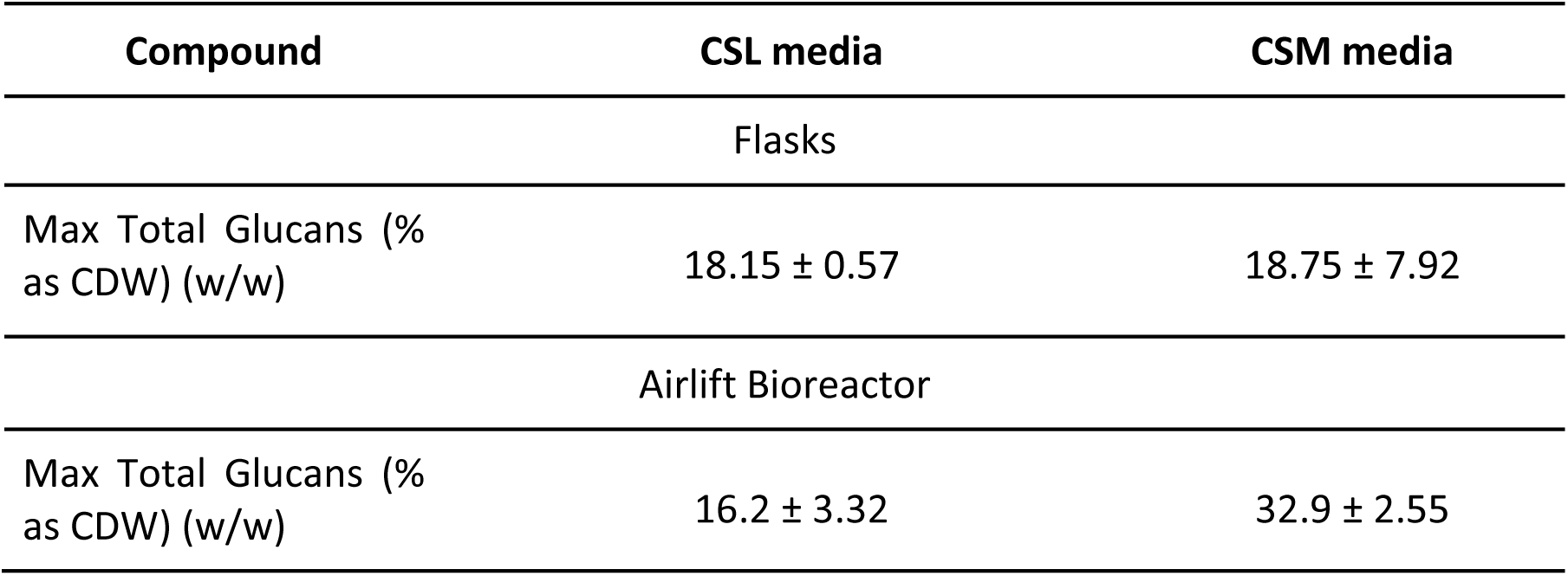
Average glucans produced by *T. versicolor* grown on spent CHOgro® Expression Media amended media (CSM) and in corn steep liquor (CSL) amended media in flasks and in airlift bioreactor. Standard deviation (SD) measurements for flask experiments had n = 3, for bioreactors n = 2. Original data values used for calculations available in table S2.

CSM was added as a 15 mL/L sterile supplement to the airlift bioreactor medium (Figure 3) and the maximum average *T. versicolor* biomass produced was 7.0 ± 0.61 g/L after 40 hours of batch growth; this value was also equal to the airlift control using CSL (Figure 3B). The volumetric productivity (as g/L/h) was higher in the airlift than in the flasks i.e.: 0.17 vs 0.037, a 4.6-fold improvement. The yields (g of biomass/g of glucose) were similar between the airlift and the flasks (i.e.: 0.7 g/g). The total glucan production in the bioreactor was 55% higher on average than in the flasks when using CSM as supplement but no difference in levels were seen between flasks and airlift bioreactor when using CSL supplement. Total glucan yield as % cell dry weight in airlift bioreactors were 2-fold higher (32.9 ± 2.55 v 16.2 ± 3.32) in cultures fed with CSM (Table 2).

**Figure 3.**
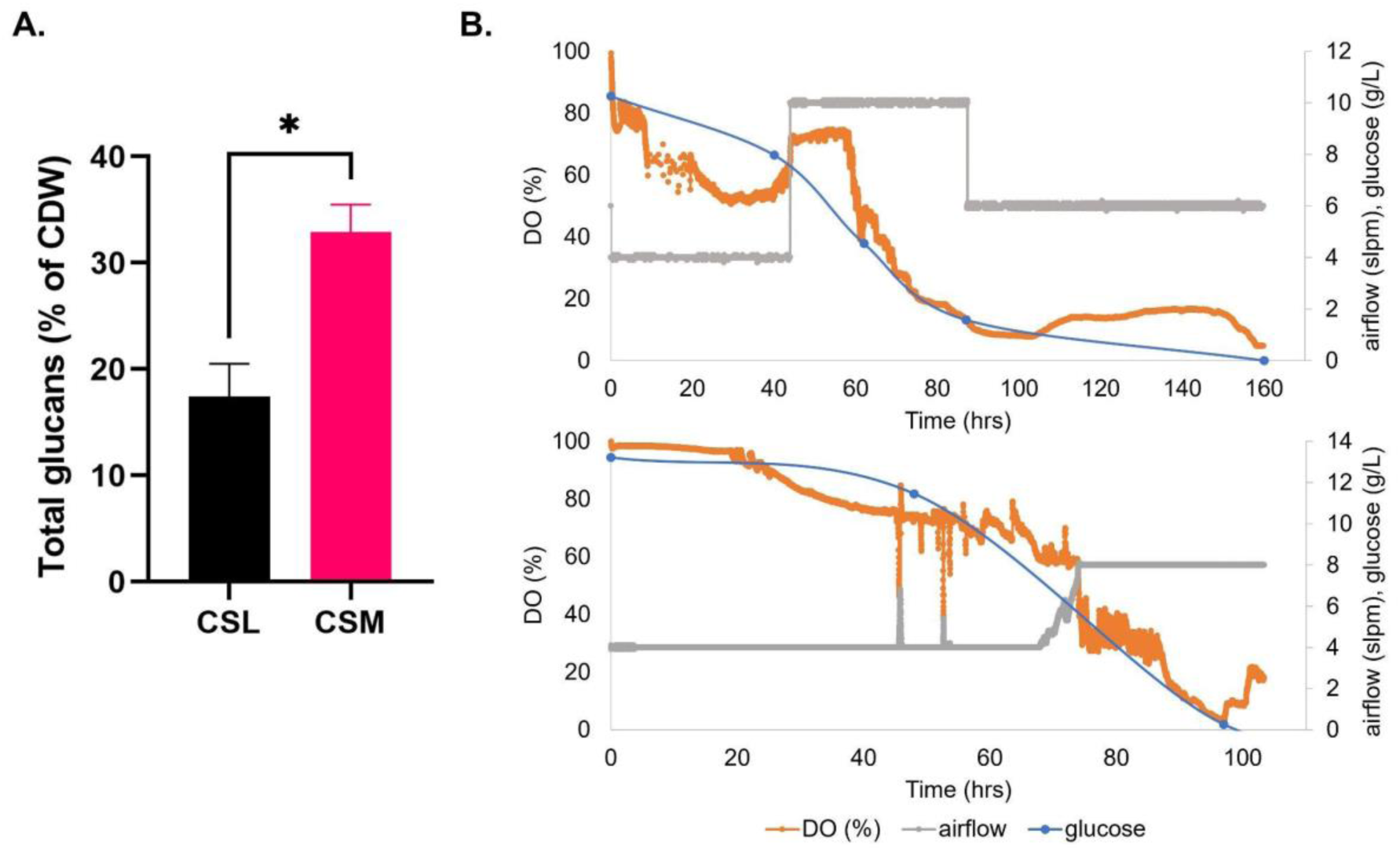
**A)** Total glucans produced by *T. versicolor* after growth on media that are CSL-based (corn steep liquor) or CSM-based (CHO spent media). **B)** Dissolved oxygen trends, carbon source consumption, and air flow over time in batch fermentation of *T. versicolor* grown on CSL-based media (top portion) or CSM-based media (bottom portion) in airlift bioreactor.

### Spent media composition analysis

Elemental analysis of the CSM and FSM revealed distinct profiles in each media with 14 of the 31 elements tested demonstrating statistically significant differences in concentration (Figure 4, panel A). Some elements were also either entirely absent or only present in trace amounts. For example, beryllium was present in the FSM but lacking in the CSM. Every other element tested for was present to some extent in both media, but to varying degrees. Manganese in the FSM (∼98 μg/L) was at significantly higher levels than in CSM (∼0.05 μg/l), while sodium was higher in the CSM (∼179 mg/) compared to FSM (4 mg/L). In most of the significantly changed 14 element levels, FSM had higher concentrations than CSM, with the exceptions of sodium and strontium.

**Figure 4:**
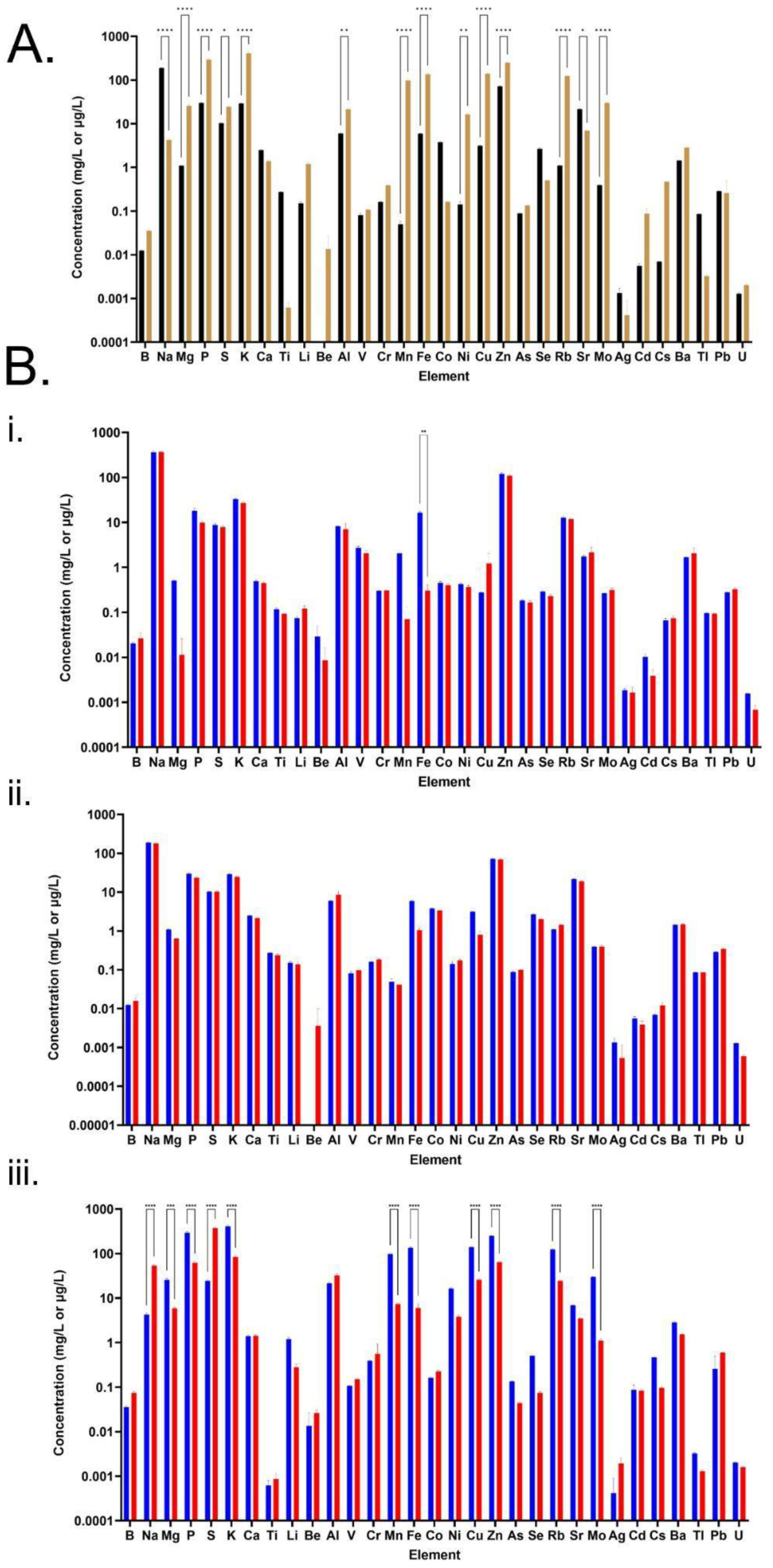
Elemental profiles of media. **A:** Concentration profiles of the two spent media prior to secondary cultures but after primary culture, depicting concentration in mg/L for elements B - Ti, and in μg/L for Li - U. Black bars = CSM post-CHO, brown bars = FSM post-fungal. Error bars represent standard deviation where n = 3 and statistical significance is indicated by asterisks after a two-way ANOVA analysis. Samples were completed in triplicate, with error bars representing standard deviation. **B:** *E. coli* consumption of elements, seen through elemental profile of media before (blue) and after (red) *E. coli* secondary flask cultures after 18 hour expression. Error bars represent standard deviation where n = 3 and statistical significance is indicated by asterisks after a two-way ANOVA analysis. **i:** LB media elemental profile. **ii:** CHO spent media (CSM) elemental profile. **iii:** Fungal spent media (FSM) elemental profile.

According to the consumption profile of all media by the *E. coli* cultures (Figure 4, panel B), the *E. coli* cultures grown in both LB and CSM did not change the elemental composition significantly with the only significant change seen in LB where iron was essentially fully consumed. The CSM condition had no significant change, though there was now an increased trace presence of beryllium after the *E. coli* culture. By contrast, the FSM composition changed significantly, with 11 elements showing significantly altered concentration levels, including two which were increased post-culture; sodium and sulphur. Consumption of potassium was ∼327 mg/L, and phosphorous was ∼232 mg/L) compared to LB and CSM fed cultures (K of ∼6 mg/L, ∼4 mg/L, and P of ∼8 mg/L, ∼6 mg/L). In addition to this increased consumption, the FSM also contained increased sulphur levels post-*E*.*coli* culture (difference of ∼350 mg/L) a higher concentration than was seen in the LB or CSM media (difference of ∼1.5 mg/L and ∼0.1 mg/L respectively). High levels of zinc were also consumed by the *E. coli* cultures grown in the FSM, with ∼186 μg/L consumed from FSM compared with ∼10.9 μg/L and ∼2.1 μg/L from LB and CSM respectively. See supplementary materials for a table of all elemental concentrations before and after *E. coli* culture (Table S3).

Amino acid composition analysis of the medias revealed that the two spent media conditions were also distinct in the profile of amino acid consumption post *E. coli* culture compared with pre-culture, and when compared to LB media (Figure 5). Both the CSM and LB conditions contained higher concentrations of amino acids than the FSM media prior to *E. coli* growth. FSM had low concentrations of amino acids left after the primary fungal culture. Interestingly, the CSM fed *E*.*coli* culture showed increased levels of some amino acids, with higher concentrations seen in amino acids alanine, cysteine, and valine. The FSM fed culture also led to a slight increase in alanine and valine post-secondary culture, but this was not statistically significant (Figure 5C). In LB fed culture there was significant consumption of amino acids glutamine, serine, lysine, aspartate, threonine, and glycine with over 1 mM of each consumed between the pre and post *E. coli* measurements. The CSM fed cultures had slight differences with over 1 mM serine, aspartate, threonine, glutamate and glycine consumed by the *E*.*coli*, and with 3 mM of serine and aspartate specifically consumed. This was double the amount of serine and almost triple the amount of aspartate consumed by the LB-fed *E*.*coli*. Synthesis or release of amino acids was also measured with 0.65 mM of cysteine and approximately 0.2 mM of alanine and valine produced. The FSM-fed *E*.*coli* had greatly reduced starting quantities of amino acids and the consumption profile was different with 0.2 mM lysine consumed. Leucine and glutamate were the second and third most consumed with 0.1 and 0.09 mM consumed respectively, and levels at less than 0.07 mM for other amino acids (Figure 5C).

**Figure 5:**
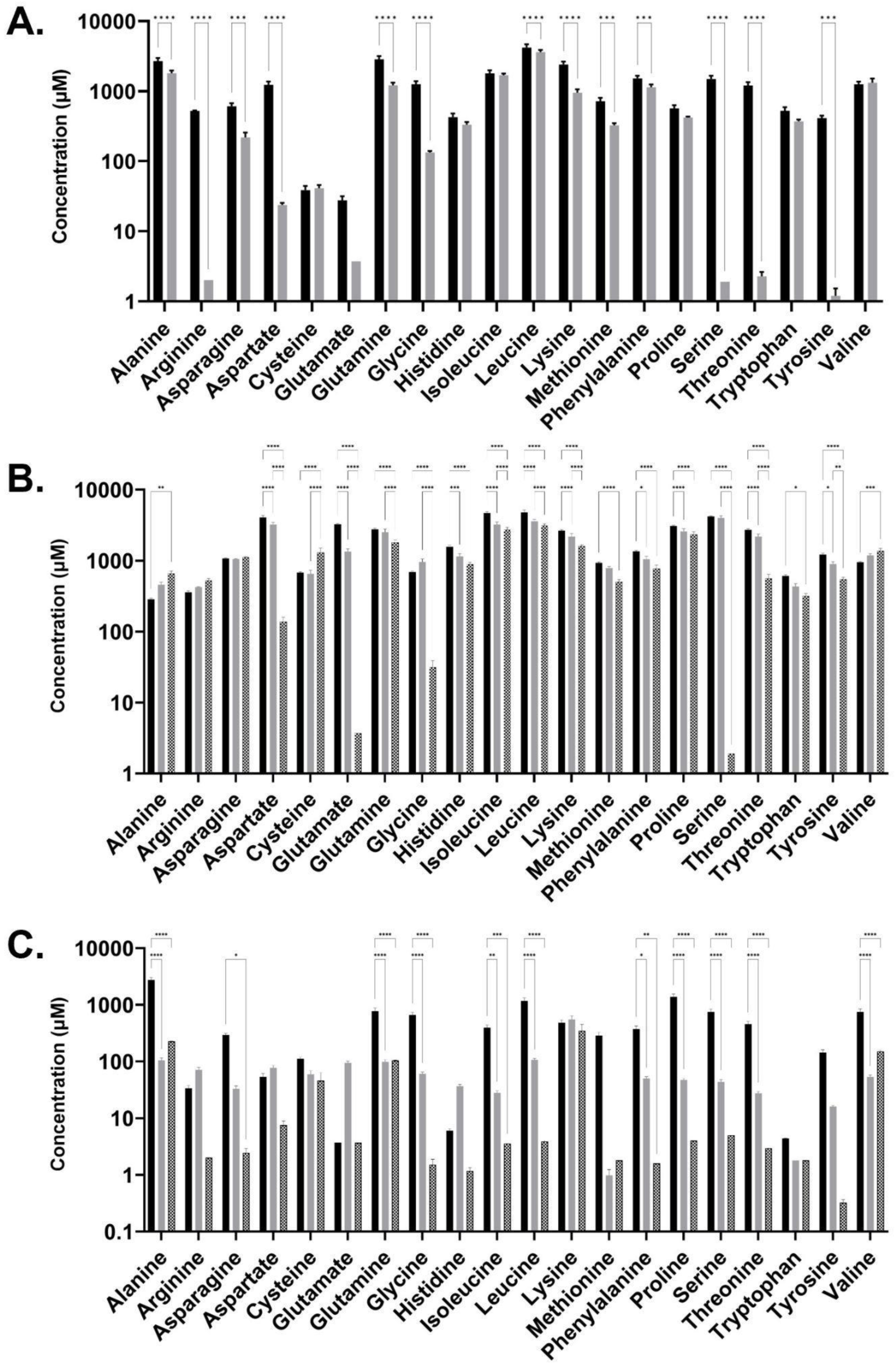
Amino acid composition of media. **A)** LB, lysogeny broth, black bars = fresh, grey = post-*E. coli* culture. B) CSM, CHO spent media, black bars = fresh, grey = post-CHO culture, hatched = post-*E. coli* culture. C) FSM, fungal spent media, black bars = fresh, grey = post-fungal culture, hatched = post-*E. coli* culture. Statistical significance was measured by two-way ANOVA and indicated by asterisks. Error bars denote standard deviation.

Analysis of carboxylic acids (Figure 6) showed that lactic acid and succinic acid were both present at varying concentrations in the media, while three other organic acids were not at high enough concentrations to affect the composition (aconitic acid, hippuric acid, and 3-Hydroxyglutaric acid). Succinic acid was produced by the *E. coli* in the CSM and LB-fed cultures though not to a significant extent, while growth in FSM did not show the same production. Lactic acid however was very high in FSM post-primary culture and was likely the reason for the low pH of this media at pH 5.5 before it was adjusted back to 7 prior to secondary culture with *E. coli*. The CHO cells also produced a large amount of lactic acid, though not to the extent of the fungal culture, and this was not consumed by the E. coli to the same degree as it was in the FSM-fed condition.

**Figure 6:**
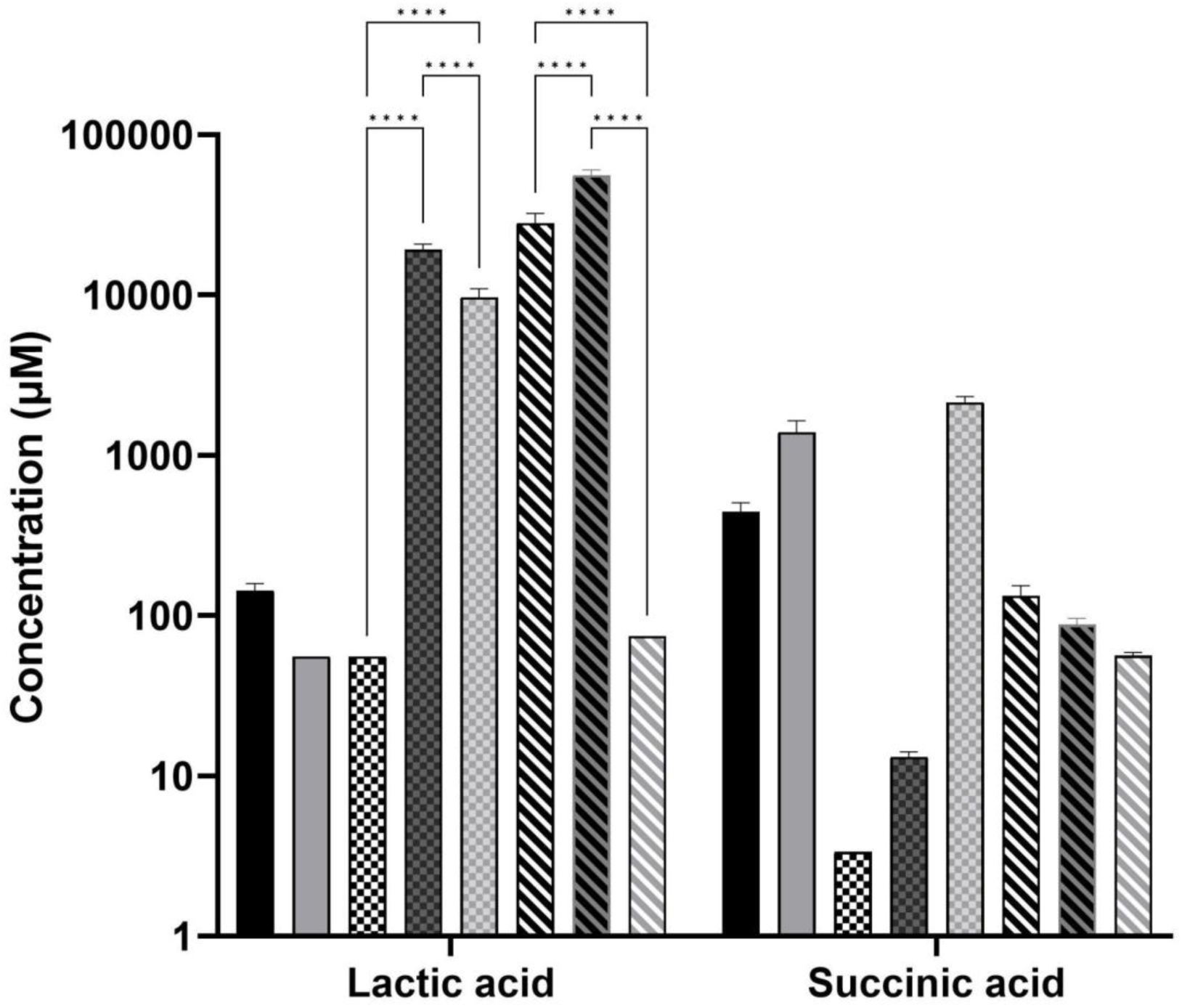
Carboxylic acid composition analysis. Error bars denote standard deviation, statistical analysis performed by two-way ANOVA with standard notation for the asterisks. Black bar = LB fresh, grey = LB post-*E.coli*, black and white check = fresh CHO media, black and dark grey check = CSM post-CHO, grey and light grey check = CSM post-*E. coli*, black and white striped = fresh fungal media, black and dark grey striped = FSM post-fungal, white and light grey striped = FSM post-*E. coli*. Other acids measured but not included due to low concentration are aconitic acid, hippuric acid, and 3-Hydroxyglutaric acid.

### Proteomic analysis of *Trametes versicolor*

Due to the increased glucan content of the CSM-fed fungal cultures, a proteomic analysis of these cultures was performed and compared to a control (CSL-fed) culture. It was found that 15 proteins were statistically significantly dysregulated after a five-day culture, and 26 proteins after a ten-day culture (Figure 7). Of particular interest were the annotated proteins of transaldolase (R7S654), phosphoglucomutase (R7S7K5), glycoside hydrolase (R7S9O5), and serine phosphatase (R7S7K2), all of which are indirectly linked to the *beta*-glucan production pathway. The first three are significantly downregulated in the CSM-supplemented condition, while the last protein, serine phosphatase is upregulated (Figure 7).

**Figure 7:**
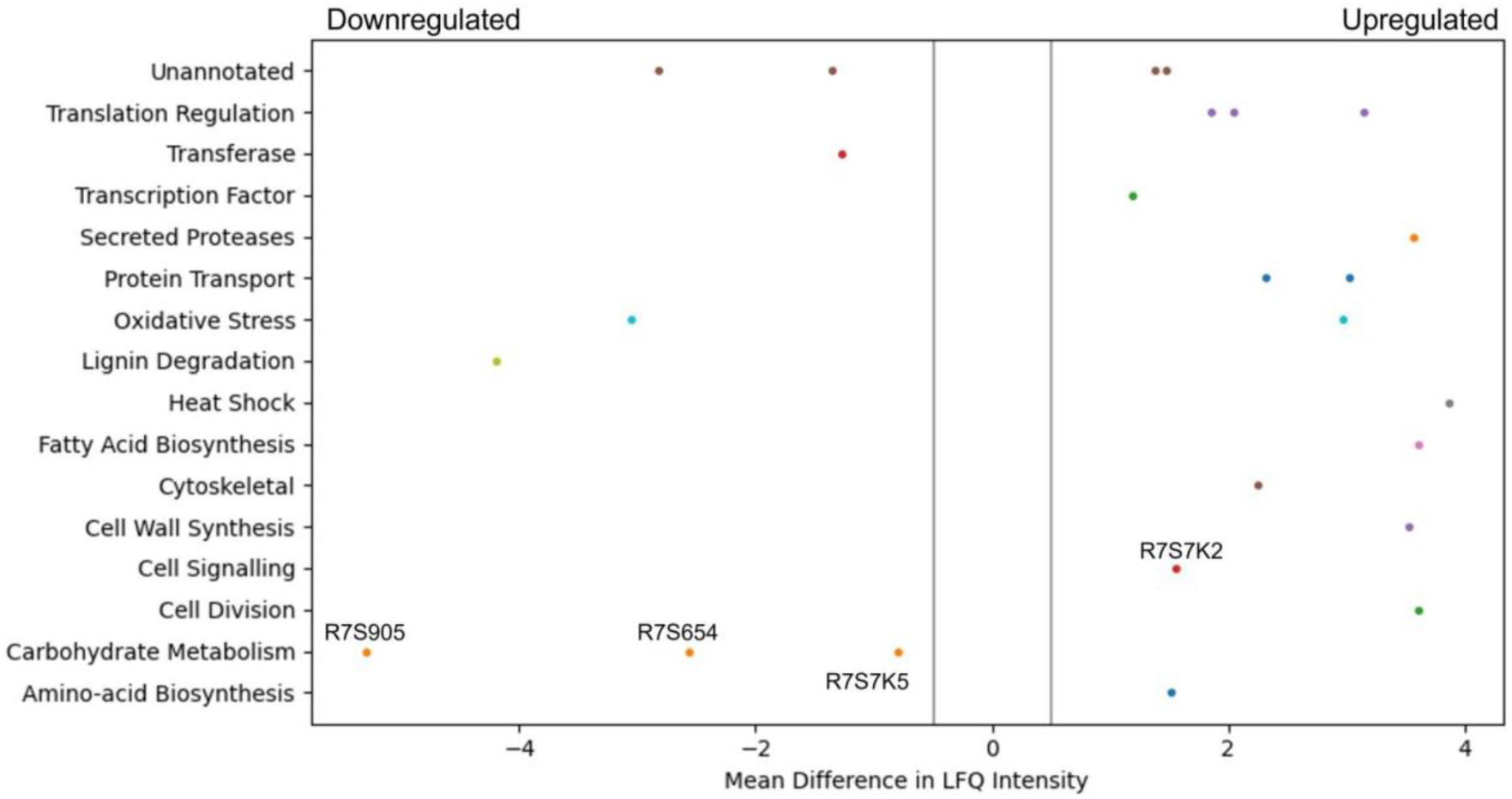
Pool Table Plot of dysregulated proteins. This graph of significantly dysregulated proteins from a 10 day culture was constructed using the Matplotlib library in Python. Significant upregulation of expression in the CSM versus CSL-fed *Trametes versicolo*r culture is shown to the right of 0 on the x-axis (−0.5 to 0.5), with significant downregulation to the left. Significance in this case was defined as having a mean difference in LFQ intensity compared to the baseline media after a student’s t-test and with an FDR of <0.05, plus a cutoff mean difference in LFQ intensity of more than 0.5, as indicated by the grey lines.

## Discussion

Cell culture media is a key component of industrial bioprocess that is immediately disposed of as waste after the primary fermentation. It is well described that many nutrients remain within media post-primary culture (Wang *et al*., 2017; O’Neill *et al*., 2022; Dodia *et al*., 2023), and this is shown in the metabolomic, proteomic and elemental analysis of the two media types used here, that we term CHO spent media (CSM) and fungal spent media (FSM) (figures 4-6).

### CHO Spent Media (CSM) supports secondary cultures

The CSM used was CHOgro® expression media, a chemically defined media used mainly for monoclonal antibody production, which is both serum and antibiotic free, making it ideal for a secondary culture feed (Voronina *et al*., 2021; Favorskaya *et al*., 2022; Rappazzo *et al*., 2022). Chemically defined media often contain the same base formulae with the addition of certain nutrients necessary for high protein expression such as L-glutamine (Ritacco, Wu and Khetan, 2018). Many of the elements identified as essential for CHO cell growth are also required by other eukaryotic and prokaryotic cells, such as magnesium, sodium, iron, and zinc (Ritacco, Wu and Khetan, 2018; Belliveau *et al*., 2021). The elemental analysis of the CSM media showed each of these elements at levels that should support the further growth and protein production in the secondary fermentations. This observation was confirmed with equivalent recombinant protein product yields in the secondary bacterial culture and a 55% increase in glucan yield seen in the CSM-supplemented *T. versicolor* fermentation (figure 3).

### Fungal Spent Media (FSM) supports a secondary culture

*Trametes versicolor* is also termed *Coriolus versicolor* and is known more colloquially as turkey tail mushroom due to a distinctive half-moon shape found fanning out from tree trunks. *T. versicolor* is a medically relevant fungus as it produces many useful molecules such as antimicrobials, including some which are secreted from its cells and have been proven to be effective against certain plant pathogens such as against the fungal species *F. langsethiae* (Parroni *et al*., 2019), and another protein which was found to have anti-leukemia properties (Ricciardi *et al*., 2017). However, no impact of antimicrobial host cell proteins (HCPs) was observed in the secondary *E. coli* culture with a specific growth rate of 88% of that for the LB condition, and equivalent recombinant protein yields at bioreactor scale as LB fed cultures.

### CSM is more efficient at feeding shake flask cultures of *E.coli* than FSM

The primary fermentation feed for *Trametes versicolor* bioprocess is a 1.5% corn steep liquor-based media (Gibbs, Seviour and Schmid, 2000). Interestingly, the FSM was found to be highly acidic (pH 5.5) after the primary culture of *T. versicolor* cultures, which had to be adjusted to 7 in order to support the *E. coli* secondary culture. This low pH was found to be as a result of the high level of lactic acid (∼56 mM) present in the media after primary culture, as shown in the metabolomic analysis of carboxylic acids (figure 6). It was also found to have significant levels of glucose (∼3 g/L) which took over 5 hours to be fully consumed by the bioreactor culture (figure 2). Though high glucose presence can inhibit the T7 expression system through catabolite repression, it was fully consumed by the induction point in the bioreactor and therefore did not inhibit the expression (Terpe, 2006). The CSM-fed secondary cultures contained even higher levels of glucose post-CHO cell primary culture (∼9 g/L), and therefore high glucose levels were not the reason for the lower protein yield seen in the FSM shake flasks. Lactic acid on the other hand is well known for its antimicrobial properties, used as protection against *E. coli* specifically in meat production (Yang *et al*., 2012). The high lactic acid levels in the FSM therefore aids in explaining the reduced protein yield obtained in the flask cultures in this media (0.216 g/L as opposed to 0.489 g/L in LB). There is precedence as it has been shown before that host cell protein (HCP) or leftover metabolite presence can affect the growth of cells in secondary cultures (Hsiao, Glatz and Glatz, 1994; Wu, Ruan and Lam, 1998; Lowrey, Armenta and Brooks, 2016; Lynch, Jordan and J O’ Connell, 2021). In the shake flask cultures of *E. coli* fed with either CHO spent media (CSM) or fungal spent media (FSM), there was a noticeable decrease in both growth rate and protein yield compared to the cells grown on nutrient rich LB media (Table 1). This is likely the result of the reduced nutrient value of the media post-primary culture, though high lactic acid concentration had an effect on the FSM-fed secondary cultures at shake flask scale. Interestingly, the bioreactor experiments with the secondary *E. coli* cultures demonstrated recombinant protein product yields that were equivalent to the rich media control condition (Figure 2). Therefore the higher dissolved oxygen, stirring, and pH maintenance at 7 is capable of restoring the protein expression performance despite the lower nutrient value and metabolite presence in the spent media feed.

The secondary culture in CSM outperformed that grown in FSM in the shake flasks, with a yield of 68% compared to 45% of the recombinant target, mCherry-EF2, compared to the LB condition. It is known that fungal cultures such as *T. versicolor* tend to excrete metabolites which can be more toxic to bacterial cultures, such as the antimicrobials mentioned before or even just more acidic metabolites in general such as lactic acid (Pereira *et al*., 2022). The reduced nutrient quantity in terms of available amino acids in the FSM post-primary culture is also a factor to be considered as this would impede protein expression (figure 5). Additionally, beta-glucans produced by the *T. versicolor* primary culture have been shown to exhibit antibacterial activity against *E. coli* at levels of over 250 mg/L (Chamidah, Hardoko and Prihanto, 2017). As *T. versicolor* is a natural producer of this polysaccharide both within its cell walls and also as a hydrosoluble compound (Vetter, 2023), any free beta-glucans that may be present within the FSM during the primary culture would have also affected the secondary *E. coli* growth, though the slightly reduced growth rate seen in the bioreactor experiments would indicate this was not a significant obstacle for the secondary culture. This finding demonstrates that a waste media containing leftover metabolites can be reused from a primary culture without any detriment to a secondary culture such as recombinant protein producing *E. coli* cultures.

### Primary carbon source choice benefitted protein production

HPLC analysis of the bioreactor experiments revealed that the leftover glucose from both spent media was completely metabolised by the secondary *E. coli* cultures after 6 hours for the CSM-fed secondary culture and after 5.5 hours for the FSM-fed culture. This aided in the quick initial growth rate in both bioreactors, at approximately 60% of that which was achieved in the LB. This remaining glucose was then completely metabolised by the protein expression induction point, a beneficial consumption point as it is known that the presence of glucose inhibits the expression of recombinant proteins using the Lac operon dependent T7 expression system (Rosano and Ceccarelli, 2014). The cultures then switched to metabolising glycerol which was added to the media at the beginning of the culture, as indicated by the slight decrease in glycerol concentration seen in figure 3. Glycerol supplementation in *E. coli* cultures has been shown to aid in maintenance of high cell density and protein production (Ganjave *et al*., 2022). While glucose is the preferred carbon source for *E. coli* growth, there are some advantages to using glycerol as the primary carbon source; Glycerol has been shown to reduce the accumulation of the growth-inhibiting metabolite acetate (Martínez-Gómez *et al*., 2012), is cheaper than glucose as a waste byproduct of the fuel industry itself, and can also reduce the sequestering of subsequently lost recombinant protein product into inclusion bodies (Murarka *et al*., 2008; Kopp *et al*., 2017).

### Beta-glucan yield was higher in secondary cultures fed with CSM

CSM supplemented cultures of *T. versicolor* grown in either shake flask or bioreactor achieved a similar biomass yield to the control corn steep liquor (CSL) supplemented cultures (Table S2). These cultures generated a 62% higher yield (g of substrate/g of biomass) than previously reported where an olive washing water substrate was used (Cerrone *et al*., 2011), with a similar yield reported by a second study in 2014, and 9% lower than reported by a more recent study in 2016 (Wang *et al*., 2014, 2016). However, the volumetric productivity of cells grown in the airlift bioreactor was 4.6-fold higher than the flask grown cultures due to both more efficient aeration and nutrient mass transfer in the airlift compared to the flasks. The volumetric productivity was between 3.36 and 5.5-fold higher than reported by the previous studies (Wang *et al*., 2014, 2016). The volumetric productivity of beta-glucans was 2-fold higher than reported by Rau et al. in 2009 and 7.5-fold higher than what was reported by Duvnjak et al. in 2016 (Rau *et al*., 2009; Duvnjak *et al*., 2016). Given that there was no inhibition of fungal growth in the flasks nor in the airlift bioreactor, the total protein concentration provided by the CSM was sufficient for the metabolic requirements of the fungal strains. Both media components were added at the same total protein concentration. Therefore, disregarding the qualitative composition of the CSL or CSM, the spectrum of the amino acids is nonetheless sufficient to guarantee the growth of the fungal strain despite one being a waste feed source.

Optimal oxygen transfer is often more vital to biomass growth than the source of protein supplement. Even more interesting was the production of bioactives by *T. versicolor* (namely, glucans); the improved biomass productivity in the airlift bioreactor resulted in a 55% increase in total glucans when CSM was used as a supplement compared with shake flask cultures (Figure 3). This increase did not occur in the CSL-based media (airlift *vs* flasks experiments). This result was believed to be as a result of the specific amino acid composition of the CSM or by some soluble molecules released by the CHO cells inducing a higher glucan production by *T. versicolor.* It is worth noting that specific amino acids could be a convenient source of glucogenic precursors that could promote the extra production of beta-glucans. One known glucogenic amino acid is serine, which was determined to be at high concentrations in the CSM media (Figure 5B). The proteomic analysis of the CSM-fed fungal cultures also showed four interesting proteins which are dysregulated in *T. versicolor* grown on CSM media; transaldolase (R7S654), glycoside hydrolase (R7S9O5), phosphoglucomutase (R7S7K5), and serine phosphatase (R7S7K2), the last of which is upregulated while the former three are downregulated (Figure 7). These proteins are known to be tied either directly or indirectly to glucan production and this led to the hypothesis that this pathway of serine-originating precursors were actively promoting beta-glucan production (Li *et al*., 2020; Santos *et al*., 2020; Tripathi *et al*., 2020; Duan *et al*., 2022). In particular, the upregulated serine/threonine phosphatase is involved in cell wall maintenance and beta-glucan production in a direct manner via cell wall-mediated stress response, and therefore partially responsible for the increase in glucans seen (Jiang *et al*., 1995; Fuchs and Mylonakis, 2009; Ariño, Velázquez and Casamayor, 2019). Succinic and lactic acid, two potentially gluconeogenic substrates, were also detected in relevant concentrations in the CSM supernatant prior to secondary culture (Figure 6). It has been shown that the regulation of the metabolic pathways involving gluconeogenesis in filamentous fungi can utilise these substrates when glucose is diminished in the media (Hynes *et al*., 2007; Chew and Than, 2021); this aids in explaining why CSM considerably increased the beta-glucan production in the secondary *T. versicolor* cultures.

### FSM was high in zinc and iron elemental composition

The elemental composition of both spent media were very distinct from one another, which is interesting in itself because both managed to successfully support the growth and protein production of subsequent cultures. The elemental composition of *E. coli* requires that certain elements be scavenged from its environment to ensure cell growth and survival (Belliveau *et al*., 2021). From the BioNumbers database (Search BioNumbers - The Database of Useful Biological Numbers (harvard.edu)), the top four elements needed from the environment for *E. coli* growth are carbon (∼50% of cell biomass), nitrogen (∼15%), phosphorus (∼3%) and sulphur (1%) with various other trace elements making up the remainder such as iron, calcium and zinc (Milo *et al*., 2010). The elemental consumptive patterns of the different *E. coli* cultures is viewable in figure 4, panel B. The much higher levels of zinc absorbed by the *E. coli* cultures grown in the FSM was a particularly interesting finding when taken with the high levels of lactic acid in the media post-primary culture. Zinc, along with iron and magnesium, has been shown to have protective properties for lactic acid-injured *E. coli* and can aid in cellular healing (Shi *et al*., 2019; Zhang, Zhang and Shi, 2022). However, high levels of intracellular zinc has also been shown to interfere with iron-sulphur cluster biogenesis, by competing for binding to iron-sulphur cluster sites on certain proteins (Li *et al*., 2019). This offers an explanation for the increased sulphur content of the FSM post-*E.coli* culture compared to the other two media where the biogenesis of sulphur containing compounds was inhibited and therefore may have increased the secretion of this element into the media.

### Amino acid consumption patterns show low precursor levels in FSM

In a previous study, it was shown that *E. coli* grown in CSM-fed cultures underwent a proteomic shift with significant upregulation of amino-acid biosynthetic enzymes (Lynch and O’Connell, 2022). This finding is reinforced by our findings here on the amino-acid content of the CSM composition pre and post-secondary culture, with the concentrations of three different amino acids (alanine, cysteine, and valine) increased after the secondary *E. coli* culture. Interestingly, alanine and valine levels were increased in both spent media conditions but not LB (FSM levels were not statistically significant). In terms of the amino acid consumption, the CSM-fed *E. coli* consumed very high concentrations of serine and aspartate with over 3 mM consumed. Both of these amino acids are precursor amino acids used to synthesise other amino acids like glycine (Pizer and Potochny, 1964; Pizer, 1965; Viola, 2001). Glutamate, another precursor, was also consumed at a high level in this condition (1.3 mM) and is one of the most essential amino acids as it has many uses outside of protein synthesis including assimilation of nitrogen as ammonium (Walker and van der Donk, 2016). Of the enzymes involved in the amino acid biosynthetic pathway found previously in CSM-fed secondary *E. coli* cultures, most were involved in the conversion of these precursors into other amino acids, including the most significantly upregulated enzyme, asparagine synthase A, with a difference in LFQ intensity of 2.79 (Lynch and O’Connell, 2022). These findings suggest that stringent response due to amino acid starvation may influence the CSM-fed *E. coli* culture in order to upregulate these biosynthetic pathways. These precursor amino acids are present at much lower concentrations in the FSM post-primary culture, having been mostly consumed by the fungal culture. This media may therefore benefit from a targeted supplementation of these precursor amino acids to improve the slightly inhibited growth rate, though the recombinant protein yields obtained were still equivalent to the LB rich media. Overall, these results lead to the conclusion that amino acid biosynthetic pathways are upregulated.

The recombinant product of mCherry-EF2 has approximately 10% of its sequence consisting of glutamine, glycine, and lysine. It is therefore unsurprising that the precursors of these amino acids (serine for glycine, glutamate for glutamine, and aspartate for lysine) are all consumed by the CSM-fed secondary cultures at high levels, as serine and aspartate were in the LB-fed condition. The secondary *E. coli* cultures fed with CSM also demonstrated an increased production of succinic acid according to the metabolomic data, though not statistically significant. A similar increase, though to a lesser extent, was seen in the rich media condition. The FSM-fed secondary cultures did not produce any measurable succinic acid. This was found to be a likely result of the high iron levels (135.65 μg/L) present in the FSM media prior to secondary culture compared to the CSM (5.92 μg/L) and LB (16.53 μg/L). Succinic acid secretion is a known response from iron starvation due to glyoxylate shunt activation in the TCA cycle (Andrews, Robinson and Rodríguez-Quiñones, 2003) particularly when coupled with glycerol as the primary carbon source (Murarka *et al*., 2008). FSM could benefit from amino acid precursor supplementation while the CSM may benefit from supplementation of certain essential elements such as iron. These supplementations may increase the growth rate, while the recombinant product yield and the fungal beta-glucan yield are already at optimal levels without further supplementation but could possibly be enhanced with such an approach.

### Conclusion

In summary, we have shown that both CSM and FSM are viable waste streams for reuse, either as a sole feed for secondary fermentations or as a protein-rich supplement. This resource has significant potential to be useful for the production of biomolecules from prokaryotic or eukaryotic organisms. The integration of primary and secondary bioprocesses with carefully selected expression hosts represents an exciting technology capable of driving a pipeline of bioproducts with a sustainable and circular model of bioproduction.

## Materials and Methods

### Mammalian cell culture

Chinese Hamster Ovary (CHO) cells were incubated in 20 mls of serum-free CHOgro® Expression media supplemented with 4 mM L-Glutamine in T75 adherent cell line flasks with filter cap, at 37°C with 5% CO_2_. Cells were split 1 in 10 once grown to 70 - 80% confluency every 3 - 4 days. Spent media was harvested at each split and was clarified of cells and cellular debris by centrifugation at 300 xg for 4 minutes prior to storing at 4°C for up to 14 days.

### Bacterial cell culture starter and transformation

Competent *E. coli* BL21 Gold were transformed by heat shock with an mCherry-EF2 fusion protein expression construct with a T7 promoter expression system and ampR, and maintained on an agar plate with 1% glucose and 100 µg/ml ampicillin. A 50 ml starter culture of these transformed E. coli was grown by incubating a single, isolated colony in Lysogeny Broth (LB) media, 2% glucose, and 100 µg/ml ampicillin. The culture was grown in a shaking incubator at 250 rpm, 37°C for 16 hours before use in the bioreactor experiments.

### Flasks-based bacterial expressions

Prior to expression, the harvested spent media, both CHO spent media (CSM) and fungal spent media (FSM), had their pH adjusted to 7. Prior to this step, the pH of the CSM was 7.2, and the pH of the FSM was 5.5. The four hour expressions were carried out in triplicate 50 ml cultures in 250 ml flasks with 2% glycerol, 100 100 µg/ml ampicillin and a 1 in 20 dilution of starter culture in the media to be tested (LB, CSM, or FSM) in a shaking incubator at 250 rpm, and 37°C. Expression of the fusion protein was induced by addition of 1 mM IPTG after the OD_600 nm_ reached 0.6. Cultures were then incubated on a shaking incubator at 250 rpm, 30°C for 18 hours. Cultures were then spun at 4°C, 4000 rcf for 20 minutes to harvest the pellets which were weighed and then frozen at −20°C prior to protein purification.

### Bioreactor-based bacterial expressions

The bioreactor used was a Bionet(R) Baby-F0 stirred tank reactor with a working volume of 1 L. The system was autoclaved prior to use. 500 ml of the media being tested was then added to the bioreactor; if a spent media was used, this was first filter sterilised using 0.22 micron vacuum filters, and 2% glycerol and 100 µg/ml ampicillin added. The LB control experiment simply had the addition of the 2% glycerol and 100 µg/ml ampicillin. System parameters used were as follows; temperature = 37 °C, pH = 7, DO = 20%, air = 0.5 slpm, and stirring = 500 rpm. A cascade was set up as well for agitation (1500 rpm max) and air (1-10 slpm), when DO drops below 20%, with agitation set up to be activated first. Acid used for pH balancing was 15% (v/v) Sulphuric Acid, and base used was 1M Sodium Hydroxide. Starter culture (50 ml) was added to the bioreactor media (500 ml) once the temperature and pH was balanced, and this was considered as the time point 0. Culture was grown until the OD_600 nm_ was > 10, with measurements taken every half an hour. Also taken every half an hour, a 1 ml sample was spun down at 14,000 rcf for 3 minutes, and the pellet kept for freeze drying in order to measure cell dry weight. 400 μl was also taken every half an hour for HPLC analysis of carbon source concentration. Once the correct OD was reached, 1mM IPTG was added to the culture with an additional 2% glycerol to maintain the cultures carbon source. From this point, an 18 hour expression was carried out with the temperature dropped to 30 °C, and left overnight. At the end of the expression, two 5 ml samples were taken for protein purification and yield measurement, by centrifuging the samples at 4000 rcf for 20 minutes. Supernatant was discarded, and the pellets were frozen at −20 °C until purification.

### Flasks-based fungal glucans production

*T.versicolor* strains were grown in 100 mL of a previously autoclaved glucose-based media in 250 mL flasks at 25 °C and 150 RPM provided with a stainless steel spring to facilitate adhesion; these are the starter cultures; these fungal strains were added aseptically into the flasks using 5 mycelia-covered agar plugs, taken from 3-days old mycelia covered malt-agar plates. The glucose based media was composed of: 10 g/L glucose, 15 mL/L of corn steep liquor (CSL) and 3.3 g/L of KH_2_PO_4_. After 5 days of growth in submerged conditions the *T. versicolor* biomass was homogenised for 30 seconds using a T18 Digital Ultraturrax(R) IKA disperser provided with S 18 N 10G dispersing tool, until the grown fungal biomass was dispersed into a homogenous population of approx 0.5 mm individual size pellets; 5mL of this homogenised biomass was aseptically transferred with a sterile 10 mL pipette into 250 mL flasks (with a 100 mL working volume) of the same media as the starter culture except that the CSL was substituted by filter-sterilised CHO spent media (CSM) supernatant. The addition of CSM was added to the flasks to have the same protein amount as the one provided by the CSL. The strain was grown in triplicate flasks. Total glucans production was evaluated by a specific enzymatic kit (K-YBGL, Megazyme(R), Ireland) assaying the lyophilised fungal biomass post-harvest after the flasks growth.

### Air-lift bioreactor-based fungal glucans production

A custom manufactured Bionet(R) double-jacketed transparent, 4L working volume air-lift bioreactor with an internal glass draught tube was used for the growth of *T. versicolor* strains. The height and the diameter of the glass vessel of the airlift bioreactor were 601 and 236 mm, respectively. The glass draught tube height and diameter were 360 and 65 mm, respectively and the draught tube is suspended inside the vessel by a double stainless-steel ring system at an approximate distance of 40 mm directly above the air sparger. The stainless steel ring air sparger has 8 1-mm holes at 0.8 cm distance from each other. The air sparger allows maximum flow-rate of 18slpm (max 4.5vvm). The design of the air-lift bioreactor (especially the height to diameter ratio) allows an internal convective movement of the sparged media with a perfect cyclical mixing approximately every 17 seconds. The media rises from inside the draught-tube due to the air sparging and sinks in the annular shape space (down-comer) externally of the draught tube but internally of the glass vessel. This gradient of media density is maintained throughout the fermentation provided that the sparging does not cease. This movement is the same that allows the pelletised biomass of *T. versicolor* to constantly recirculate in the airlift bioreactor. Aliquots of pelletised biomass were aseptically sampled at different time intervals from the airlift bioreactor, using a sterile withdrawing 16mm wide silicon tube and an external wide head peristaltic pump. These biomass aliquots were frozen at −70C and then freeze-dried and used for glucans and cell dry weight analyses (CDW). The bioreactor was used in batch mode (i.e.: all the media components are added at the beginning of the fermentation); the media composition was the same as the one in the flasks experiments except that CSL was replaced by filter sterilised CHO cell spent supernatant; the media composition was the following i.e.: 10 g/L glucose, 15 mL/L of filter sterilised spent CHO cell supernatant and 3.3 g/L of KH_2_PO_4_. The inocula was taken from a 2 L flask where the *T. versicolor* pelletised biomass was grown for 72 hours from a homogenised starter culture. The *T. versicolor* pelletised biomass was homogenised by an Ultraturrax for 30 seconds at 18000 RPM in aseptic conditions (in a SafefastTM Class II laminar flow cabinet) and added to an empty autoclaved Duran bottle provided with sterile silicone tubing so to be added to the airlift in aseptic conditions.

### Specific growth rate calculations

Specific growth rate constant (k) and doubling time was calculated by use of GraphPad Prism’s Exponential (Malthusian) growth equation, version 9.4.1 for Windows, GraphPad Software, San Diego, California USA, www.graphpad.com. Plots were generated from growth curves of OD_600 nm_ versus time (hrs), with triplicate values.

### Recombinant protein purification and yield calculations

Cell pellets were resuspended using a buffer containing 10 mM Tris and 2 mM CaCl2 pH 8 before lysis by sonication. The lysate was then boiled at 85 °C for one minute and spun at 15,000 rcf for 30 minutes. Supernatant was harvested and referred to as the post-boil (PB) sample. Protein concentration was then measured with a DeNovix DS-11 Spectrophotometer, using the UV-Vis application. Measurements were taken at 585 nm for mCherry yield, with an extinction coefficient calculated to be 44,854.2 M^cm−1^ and a protein molecular weight of 30,667.4 g/mol. Total yield was then calculated using Beer-Lambert’s law.

### High Pressure Liquid Chromatography (HPLC)

The analyses for glucose/glycerol concentrations in the media supernatant were performed using an HPLC unit. This analysis was carried out on a Shimadzu Prominence unit (SIL-20AC HT autosampler, DGU-20A5 degasser, CTO-20A column oven, RID-10A detector) fitted with a Bio-Rad Aminex HPX-87H ion exclusion column. The method used is an isocratic elution with a mobile phase of 0.014N sulphuric acid. Flow rate is set at 0.55 ml/min with a pressure set point of 45 kfg (4.41 MPa). The compounds are detected by a RID (refractive index detector). The RID cell is equilibrated and balanced between each sample injection. Dilutions of the media supernatants were used in Whatman(R) MIni-UniPrepTM syringeless filter vials and 20 uL of the supernatant was injected into the HPLC column from each vial. Standard curves of the aqueous solutions of analytical grade glucose or glycerol were constructed using the same methodology and the dilutions of the supernatant media samples were only considered when they fell into the concentration ranges of the standards.

### Inductively Coupled Plasma - Mass Spectrometry (ICP-MS)

Media samples for ICP-MS were collected by harvesting 1 ml sample volumes in triplicate. Fresh media samples were prepared and harvested immediately, of fresh LB, fresh Corn Steep Liquor (for fungal growth) and fresh CHOgro® expression media (for CHO cell growth) all in triplicate. Fungal spent media (FSM) post fungal culture was harvested after a 72 hour culture of *Trametes versicolor* in the airlift bioreactor as above. CHO spent media (CSM) samples were harvested from a three day culture of adherent CHO cell culture. LB, CSM, and FSM post *E. coli* cultures were harvested after the 18 hour expression experiment carried out in shake flask culture. All samples were diluted 1 in 10 with 2% nitric acid in milliQ water, and sent to University of Nottingham for ICP-MS analysis.

## Acknowledgments

CL is funded by the Atoms-2-Products center for doctoral training, which is supported by the Science Foundation Ireland (SFI) and the Engineering and Physical Sciences Research Council (EPSRC) under Grant No. 18/EPSRC-CDT/3582. Our thanks to Katalin Kovacs, Peter License, and the ICP-MS suite at the University of Nottingham for their assistance in the analysis of our media composition by ICP-MS. Also to Lorraine Brennan and Xiaofei Yin at the Core Facilities in the Conway Institute for their assistance in the metabolomic analysis of the media supernatant.

